# Phages rarely encode antibiotic resistance genes: a cautionary tale for virome analyses

**DOI:** 10.1101/053025

**Authors:** François Enault, Arnaud Briet, Léa Bouteille, Simon Roux, Matthew B. Sullivan, Marie-Agnès Petit

## Abstract

Antibiotic resistance genes (ARG) are pervasive in gut microbiota, but it remains unclear how often ARG are transferred, particularly to pathogens. Traditionally, ARG spread is attributed to horizontal transfer mediated either by DNA transformation, bacterial conjugation or generalized transduction. However, recent viral metagenome (virome) analyses suggest that ARG are frequently carried by phages, which is inconsistent with the traditional view that phage genomes rarely encode ARG. Here we used exploratory and conservative bioinformatic strategies found in the literature to detect ARG in phage genomes, and experimentally assessed a subset of ARG predicted using exploratory thresholds. ARG abundances in 1,181 phage genomes were vastly over-estimated using exploratory thresholds (421 predicted vs 2 known), due to low similarities and matches to protein unrelated to antibiotic resistance. Consistent with this, 4 ARG predicted using exploratory thresholds were experimentally evaluated and failed to confer antibiotic resistance in *Escherichia coli*. Re-analysis of available human-or mouse-associated viromes for ARG and their genomic context suggested that *bona fide* ARG attributed to phages in viromes were previously over-estimated. These findings provide guidance for documentation of ARG in viromes, and re-assert that ARG are rarely encoded in phages.

## Introduction

Antibiotic resistance is widely recognized as problematic for the treatment of infectious diseases, due to the emergence of antibiotic resistant pathogens (World Health Organization 2014). However, antibiotic resistance genes (ARG) are also common outside of pathogens. Among 6179 sequenced microbial genomes, 84% have at least one ARG (Gibson et al 2015) including many diverse, mostly non-pathogenic bacteria in soils (Allen et al 2009, Forsberg et al 2012, Riesenfeld et al 2004, Torres-Cortes et al 2011), activated sludge (Mori et al 2008, Parsley et al 2010), as well as in humans and animals (Kazimierczak et al 2009, Penders et al 2013, Sommer et al 2009, Wichmann et al 2014). Further, ARG can be transferred between soil bacteria and human pathogens via horizontal gene transfer (Forsberg et al 2012), such that non-pathogens represent a reservoir of ARG to pathogens (Canica et al 2015, Martinez et al 2015, van Schaik 2015). It becomes critical then to estimate how often ARG are present on mobile genetic elements (Martinez et al 2015), and to understand the conditions under which these elements promote horizontal gene transfer in nature, and lead to the emergence of antibiotic resistance among pathogens.

Numerous mobile elements are implicated in the spread of ARG (Broaders et al 2013, Huddleston 2014). These include plasmids and integrative conjugative elements (ICE), via conjugation (for reviews see (Davies and Davies 2010, Wozniak and Waldor 2010)), as well as generalized transduction carried out by bacterial viruses, i.e. phages (Balcazar 2014, Davies and Davies 2010, Muniesa et al 2013, Volkova et al 2014). Quantitatively, laboratory experiments with phage P1 suggest that ARG transfer is 1000-fold less common via phage transduction than via conjugative elements (Volkova et al 2014). This is due to the fact that, contrary to conjugation events which will systematically transfer ARG together with the ICE or plasmid genome, generalized transduction relies on erroneous encapsidation of non-phage DNA. Measurements from phage P1 suggest that this is a rare event as only about 4 out of 10^4^ phage capsids will encode a chromosomally-encoded ARG gene (Volkova et al 2014). In addition, ARG are only rarely directly encoded in phage genomes - 2 of 1181 publicly available phage genomes contain an ARG (NCBI RefSeq as of April 2014, Pruitt et al 2007). These include the twin phages Gamma and Cherry of *Bacillus anthracis*, which encode a fosfomycin resistance gene (Schuch and Fischetti 2006). Beyond these, some *Staphylococcus aureus* satellite phages, which require another phage to propagate, carry ARG (Novick et al 2010). Thus, while phage-encoded ARG would presumably provide advantage to bacteria hosting the phage in a lysogenic state, there appears to be little selection for phages to carry such genes.

In contrast, recent virome studies have reported high levels of ARG in phages from pulmonary human samples of cystic fibrosis patients (Fancello et al 2011, Rolain et al 2011), and from feces samples of antibiotic treated mice (Modi et al 2013). These studies call into question how prevalent ARG are among phages and suggest that phages play far greater roles in spreading ARG. The higher ARG frequencies in viromes could derive either from genome-sequenced phage isolates misrepresenting naturally-occurring ARG frequencies, or virome studies over-estimating ARG frequencies.

The first step to identify ARG in virome sequences involves a homology search against a database dedicated to ARG. The Antibiotic Resistance Database (ARDB, 7828 proteins at last update in 2009, Liu and Pop 2009) is commonly used, sometimes together with proteins annotated with the GO term “antibiotic catabolic process” (Modi et al 2013). The use of ARDB necessitates caution, as recently observed (van Schaik 2015), and downstream manual inspection to refine *in silico* functional assignments is required. More recently, expert curated databases have become available including the Comprehensive Antibiotic Resistance Database (CARD, McArthur et al 2013) and Arg-annot (Gupta et al 2014), which are restricted to experimentally-confirmed proteins conferring antibiotic resistance, and Resfams (Gibson et al 2015), which updates CARD with recently discovered beta-lactamases. The first two contain 2822 and 1689 proteins, respectively, while the third contains 2097 proteins grouped into 166 families, using an approach similar to Pfam (Bateman et al 2000). Of these Resfam families, 119 are “core” and can be used for ARG discovery without ambiguity, while 47 are more challenging to interpret as they will also match non-ARG proteins, such as ABC transporters and transcriptional regulators.

Once a reference database is selected, a relevant sequence similarity threshold to identify ARG among a collection of new sequences must be chosen. The literature offers both conservative and exploratory options. Conservative criteria require an unknown ORF to match the database with either >40% coverage over the target ARG and >80% nucleotide identity (Zankari et al 2012), or >85% coverage and >80% amino-acid identity (Gibson et al 2015). These stringent criteria will largely only identify known ARG (Gibson et al 2015, Zankari et al 2012). However, in the last decade many new ARG have been discovered by functional screening, which would not have been found using stringent comparisons to databases (Moore et al 2013, Parsley et al 2010, Sommer et al 2009, Wichmann et al 2014). To discover distantly related ARG, one may wish to lower down similarity cut-offs, a common practice for virome studies, where high levels of sequence divergence are routinely observed, due to the high mutation rates of phages and lack of explored ‘sequence space’ resulting in limited reference genomes. In this case, E-value thresholds below 10^−5^ or 10^−3^ have been used (Modi et al 2013, Willner et al 2009).

Here we compare approaches to detect ARG in phage genomes, experimentally evaluate 4 predicted ARGs, and assess the impact of conservative and exploratory thresholds for inferring ARG from 25 published viromes, including those with high reported levels of ARG (Modi et al 2013, Willner et al 2009). We build a case that *bona fide* ARG frequencies are vastly over-estimated in virome studies, and suggest that the main path for ARG dissemination by phages is generalized transduction as commonly asserted.

## Material and methods

### Databases of ARG

The four databases compared in this analysis are an updated version of ARDB ((http://ardb.cbcb.umd.edu/, named hereafter ARDB+, 13,453 different proteins, see details below), Arg-annot (http://www.mediterranee-infection.com/article.php?laref=282&titer=arg-annot, download May 2015), CARD (http://arpcard.mcmaster.ca/download, subset download excluding genes that confer resistance via specific mutations, download May 2015), and Resfams (http://www.dantaslab.org/resfams, v1.2, updated 2015-01-27). The search against Resfams is not performed with BLAST but hmmscan (Finn et al 2011), using the–cut_ga parameter that sets the threshold for similarity according to the threshold chosen to aggregate the members of each family. ARDB+ contains the 7828 initial proteins from ARDB, complemented with 5625 Uniprot proteins (Sept. 2014) filtered for the GO:0017001 term «antibiotic catabolic process», and sharing less than 98% identity with ARDB proteins. Remarkably, 80% of these additional proteins are beta-lactamases. We observed *a posteriori* that a partition protein (whose function is to stabilize plasmids and temperate phages replicating autonomously) has been mistakenly incorporated into ARDB (XP_002333050), with the annotation “tetracycline resistance gene from *Populus trichocarpa”*. Concerning the CARP database, a thymidylate synthase from Enterococcus faecalis (AF028811.1) was similarly removed from this analysis. All results with ARDB+ and CARD are given after removal of these two erroneous matches. ARDB+ being significantly more populated than CARD, Arg-annot and Resfams, we compared the content of these databases. Approximately 88% of ARDB+ proteins were similar (with BLAST and bit-score >70) to those of CARD and Arg-annot (Supplementary information S1), and 62% only are found in Resfams (with hmmscan and the stringent built-in threshold).

### Reference set of proteins from complete phage genomes, and generation of mock viromes

Phage proteins from the 1,181 genomes (121,505 proteins) were downloaded from the NCBI viral genome database (http://www.ncbi.nlm.nih.gov/genomes/GenomesGroup.cgi?opt=virus&taxid=10239&host=bacteria, 08/04/2014). To generate the mock200 and mock580 reads from this set of genomes, successive DNA fragments of identical sizes were generated sequentially from the genome files.

### Viral metagenomic data

All viromes composed of at least 50,000 reads available at the beginning of the analysis (2014), and originating from human-or mice-associated bacterial ecosystems were downloaded and studied (25 viromes in total). Of these, 23 were available as unassembled reads: 10 viromes from human lung (Willner et al 2009), 9 from human feces (5 from Kim et al 2011); 4 from Minot et al 2011) and 4 from mice feces (Modi et al 2013). In addition, 2 already assembled datasets, both from human feces (Minot et al 2012, Reyes et al 2010), were also considered. All unassembled viromes were sequenced using different generations of the pyrosequencing technology (454, Roche) and the average read length of individual datasets is comprised between 205 and 873.

To normalize read length among viromes and have comparable results for all viromes, comparisons with the three databases were also performed on viromes where each read was randomly truncated to 200 bp.

For the mice viromes (Modi et al 2013), low-quality ends of the reads were trimmed (quality score < 20) and reads shorter than 100 bp were removed.

### Analysis of raw reads for 16S content and bacteria-only COGs

All reads from the 23 unassembled viromes were truncated to 200 bp and compared (BLASTn, bit-score>200) to the SILVA database (Quast et al 2013) for the identification of 16S rDNA. To refine bacterial DNA detection, the protein families (COGs) the most frequently observed in human digestive tract microbiomes were taken from the Qin et al. gut metagenomic analysis (Qin et al 2010). The 1,112 prevalent COGs in all individuals of their study were compared to the 172,774 proteins of all sequenced viruses present in the RefSeq database (Pruitt et al 2007) (http://www.ncbi.nlm.nih.gov/genomes/GenomesGroup.cgi?taxid=10239). Using a threshold of 50 on the bit-score, the 688 COGs having no similarities with viral proteins were further used and termed “bacteria-only COGs”.

To estimate the amounts of bacteria-only COG genes in bacterial genomes, 1,102 completely sequenced genomes from the KEGG database (version of 2011) were chopped in 580bp-long reads and compared to the bacteria-only COGs (BLASTx bit-score>50). 21.1% of bacterial reads had a hit against these COGs. To estimate the amount of ARG in bacterial genomes recovered with the bit-score 70 threshold, the same 580bp-long bacterial mock reads were compared with BLASTx against ARDB+: 2.2% of these reads had a hit against ARDB+.

### Assembly and analysis of contigs

All viromes were assembled de novo using Newbler 2.6 (454 Life Sciences), using the threshold of 98% identity on 35 bp. For each contig longer than 500 bp of the 25 assembled viromes (23 assembled in-house, 2 already assembled), an ORF prediction was processed using MetaGeneAnnotator (Noguchi et al 2008) with default parameters. Predicted ORFs were compared to ARDB+ proteins (bit-score > 70) using BLASTp, and to Resfams using hmmer.

### Cloning and testing of four putative phage-encoded ARG

**Strains, plasmid construction and testing are described in Supplementary information S2**.

## Results

### Assessing informatic stringencies to identify candidate ARG in phage genomes

To evaluate relevant databases and thresholds for identifying known ARG, we examined the number of ARG we could detect among the 121,506 proteins (termed ‘proteome’) from 1,181 reference phage genomes. As described above, only 2 ARG were expected with this dataset - the *fos* genes of phages Gamma and Cherry (Schuch and Fischetti 2006). Using a conservative threshold (>40% coverage and >80% amino-acid identity), BLASTp screens of the reference phage proteome identified the 2 positive control *fos* genes, as well as a possible beta-lactamase of phage G against ARDB+, and no hits against CARD and Arg-annot (Figure 1A; Supplementary information S3). Using the same proteome and the built-in conservative threshold of Resfams identified one hit in the *yokD* gene, an aminoglycoside acetyl transferase of *Bacillus subtilis* prophage SPbeta. Neither the phage G ‘beta-lactamase’ gene nor the SPbeta *yokD* gene are proven to function as ARG, but represent strong candidates for future experimentation. The lack of detection of the *fos* genes outside the ARDB+ database was due to the fact that the *fos* genes present in the other databases were too divergent (57-63% identity). In the following analyses, the most recent Resfams tool was used to detect ARG among protein datasets, but as Resfam is not adapted to raw DNA reads, BLASTx comparisons against ARDB+ with cutoffs of >40% coverage and >80% amino-acid identity will be used instead.

**Figure 1:**
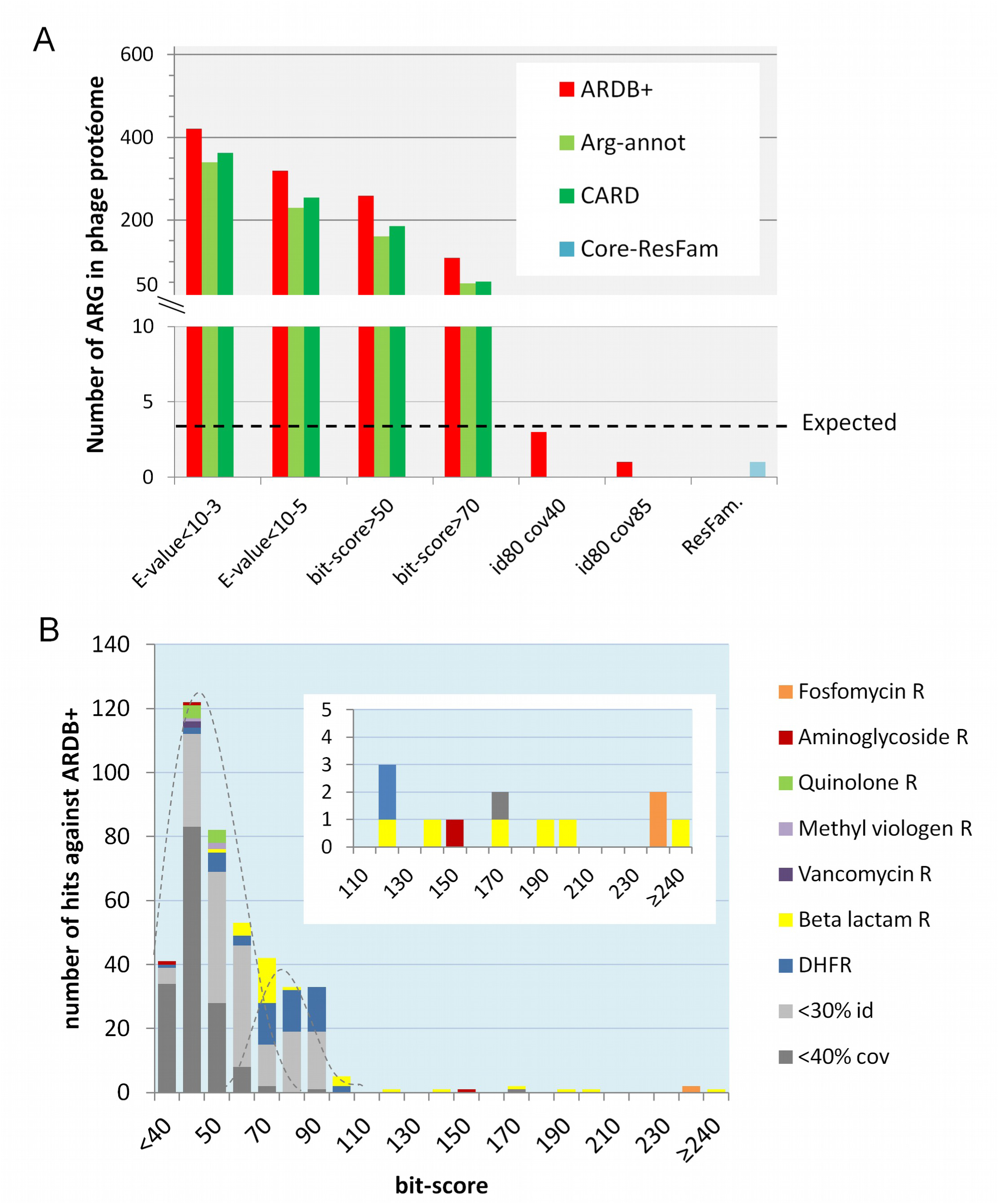
Analysis of the reference phage proteome against various antibiotic resistance genes database. **A**. Comparison of the recovery of ARG hits with the four ARG databases, using the conservative thresholds of Resfams, >40% coverage and >80% amino-acid identity, or >85% coverage and >80% amino-acid identity, as well as the exploratory thresholds E-values 10^−3^ and 10^−5^. However, as E-values on large databases cannot be simply transposed for the much smaller ARG databases, two additional bit-score thresholds (a statistics independent of the database size) were introduced, 50 and 70. The dotted line indicates the expected number of hits (2), according to experimental data. **B**. The 421 hits against ARDB+, obtained with the most exploratory cut-off (E-val < 10^−3^) are displayed, as a function of their bit-score value, with a color code for resistance category. In grey, the hits with <30% identity or <40% coverage, which are most likely false-positives. Zoom inset: hits with bit-scores > 110.

In contrast to these conservative threshold results, the exploratory thresholds resulted in hundreds of hits for all databases considered (Supplementary information S3). To better understand whether these should reasonably be considered candidate ARG, the 421 hits recovered with the exploratory cut-off (E-value<10^−3^) against ARDB+ were further examined. The bit-score distribution of these hits was bimodal with a bit-score cut-off of ~70 at the junction (Figure 1B), suggesting a first population (<70 bit-scores) of random hits, distinct from a second population (>70 bit-scores) of significant hits. We therefore manually inspected the 109 hits with bit-scores >70. Ninety-six of these could be rejected as non-significant because the target ARG was poorly covered (<40%), too highly divergent (<30% identity), or else had homology to enzymes likely to serve non-ARG functions in phages. On the latter, homology with di-hydrofolate reductase was rejected without further consideration, as this enzyme potentially conferring trimethoprim resistance is more likely to be involved in nucleotide metabolism in phages (Asare et al 2015b). Beta-lactamases were also further considered, as genes with homology to beta-lactamases likely play different roles in phages (Asare et al 2015a, Quiros et al 2014), particularly when of atypical length. Indeed, one such protein annotated as a beta-lactamase is in fact a tail fiber protein (Cresawn et al 2015). Half of the putative beta-lactamases were rejected because of atypical lengths and/or similarity to tail proteins (see Supplementary information S4 for the complete analysis of beta-lactamase hits, and Supplementary information S5 for examples of rejected hits). In total, 13 hits were retained after manual inspection, including the 2 experimentally proven *fos* genes, and 11 additional candidates that are likely worth experimental follow-up, including the aminoglycoside acetyl transferase from the *Bacillus subtilis* SPbeta prophage, and 10 beta-lactamases.

Together these results suggest that findings using the conservative threshold, even against the permissive ARDB+ database, will recover only *bona fide* ARG, whereas the exploratory thresholds may lead to the discovery of novel ARG but do so at the expense of hundreds of false-positives.

### Experimental testing of four putative phage-encoded ARG

To test whether the stringency of this manual screening, which removed 90% of the hits, was appropriate, four ARG candidates were chosen for experimental evaluation. Specifically, *yokD* of phage SPbeta, encoding a putative aminoglycoside transferase, and three beta-lactamases related to those listed in Supplementary information S4 [*gp34* of phage Palmer (99% identical amino-acids to phage Pony Gp33), *gp62* of mycophage Corndog and *gp20* of Mozy (97% identical amino-acids to *gp20* of phage Che8)] were examined. These last two putative beta-lactamases were rejected upon manual inspection, and suspected to be rather phage tail proteins (Cresawn et al 2015). All four proteins were expressed in *Escherichia coli* (see Supplementary information S2 for the origin of genes, and cloning details) and found to be soluble (Supplementary information S6). But none of the four proteins conferred antibiotic resistance, in antibiogram assays (Figure 2A). Since none of the putative beta-lactamases contained predictable signal peptides, and to exclude the possibility that heterologous expression prevented resistance *in vivo*, the activity of crude extracts was tested *in vitro*. For this, 40 μL of a soluble total protein extract was spotted on a lawn of top-agar containing an indicative *E. coli* strain which growth was prevented by the supplementation of ampicillin (or kanamycin). Inactivation of the antibiotic around the spot containing the modifying enzyme permitted local growth of *E. coli*, as shown with the positive controls (strain expressing the pBR322 encoded beta-lactamase, or the pET9 encoded aminoglycoside acetyl transferase). Again, none of the crude extracts with phage-encoded candidate ARG permitted antibiotic degradation (Figure 2B). In addition to ampicillin, the putative beta-lactamases were tested on mecillinam (another penicillin, against which a prophage gene similar to Palmer gp34 has reported activity, Ogilvie et al 2013), ceftazidime (cephalosporin), and aztreonam (monobactam) with the same negative result. We conclude that among these 4 phage genes, none was a *bona fide* ARG. The absence of resistance with the YokD protein was surprising, given its very good similarity scorings (bit-score=156, E-val=10^−46^). However, the closely related putative aminoglycoside transferase BA2930 of *Bacillus anthracis* is also unable to degrade kanamycin in antibiogram assays (Klimecka et al 2011). It is a proven acetyl transferase, but seems to have a substrate distinct from aminoglycoside antibiotics. We conclude that exploratory thresholds, even after manual curation, lead to over-estimations of ARG counts.

**Figure 2:**
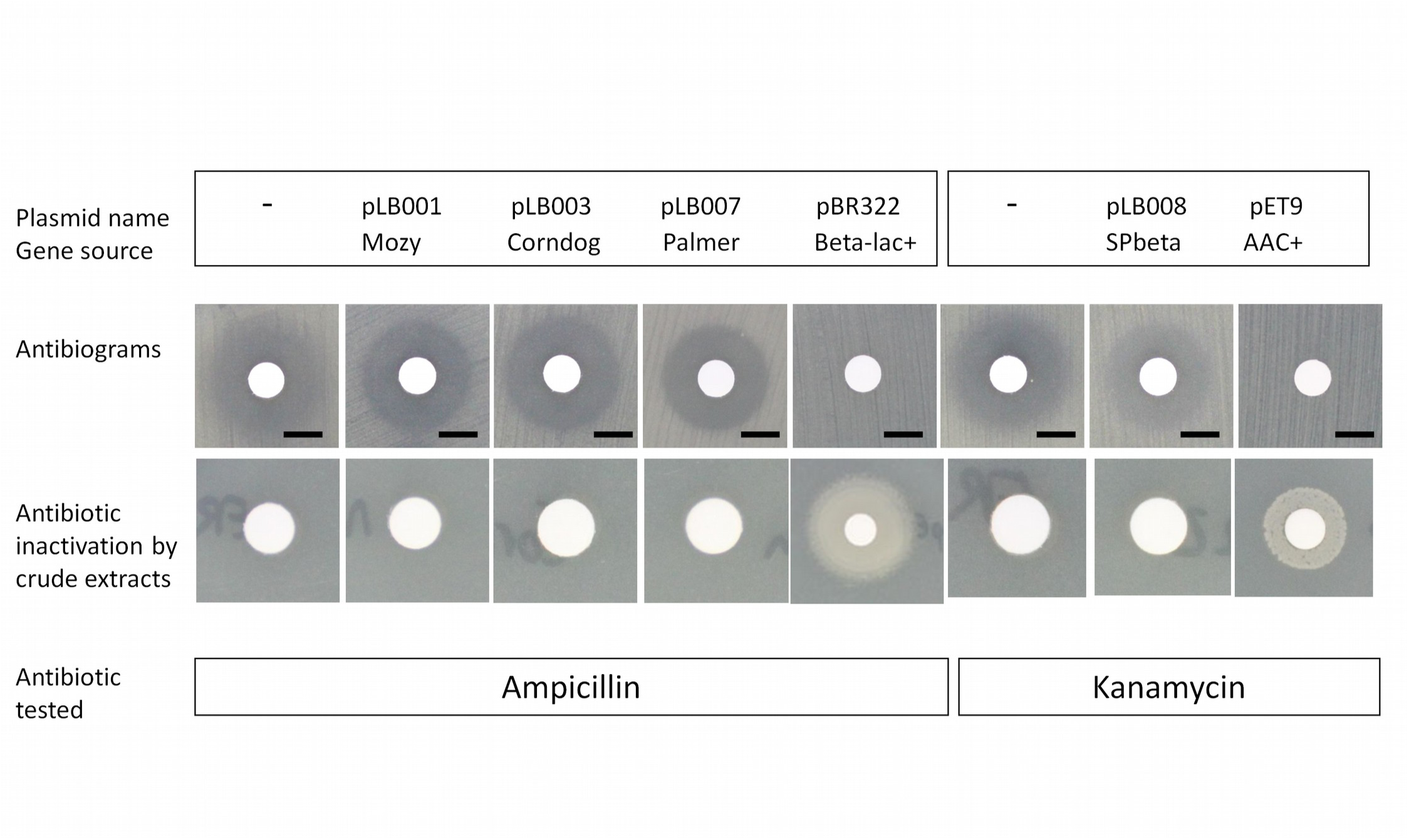
Experimental testing of four predicted ARG. A. Antibiograms. The three beta-lactamases were tested *in vivo* by spreading bacterial lawns of *E. coli* expressing each protein (100 μM IPTG) into top-agar, and spotting 10 μg of ampicillin on 6 mm diameter disks. For YokD, 30 μg kanamycin was used. Plates were incubated at 30°C. Confluent growth was observed for the pBR332 (Beta-lac+: beta-lactamase postive) or pET9 (aac+: aminoglycoside acetyl transferase positive) positive controls, but inhibition zones were present for all phage-encoded putative enzymes. Bar, 6 mm. **B. Enzymatic tests**. Ampicillin (100 μg/mL) or kanamycin (50 μg/mL), together with an indicator ER2566 *E. coli* strain sensitive to both antibiotics, were spread into top-agar. Soluble fractions of total extracts of *E. coli* ER2566 expressing each of the four proteins, or expressing the ARG of plasmid controls, were then spotted on 6 mm diameter disks. Plates were incubated for 24h at 30°C. Bacterial growth around the disk indicates that the protein extract contains antibiotic resistance activity, which degraded sufficient amounts of antibiotic in the top agar to permit growth of the antibiotic sensititive *E. coli* strain.

### Detecting ARG in virome reads

Given the above informatics benchmarking, and its conflict with experimental tests, we next sought to quantify the number of ARG in viromes. However, as opposed to the full-length proteins examined above, the length of virome reads is also critical to evaluate, as they impact ARG discovery. To this end, we mimicked currently available virome read lengths by *in silico* fragmenting the 1181 reference phage genomes to create 200 bp or 580 bp reads. These mock community viromes were termed mock200 and mock580, respectively, and then compared to the same databases (BLASTx) using the conservative and exploratory thresholds (see Supplementary information S3). Results paralleled those from the full-length proteome analyses for conservative thresholds. For the exploratory threshold based on BLAST bit-score >70, hits were fewer among short reads (0.09% of all reads for mock200 and 0.60% for mock580, against ARDB+) than among full-length proteome analyses (0.81%), so that specificity increased slightly (Supplementary information S7). Therefore, conclusions drawn from full genomes hold mostly for short reads, and suggest the use of the >70 bit-score threshold for exploratory searches, to minimize false positives.

### More ARG in viromes than in reference phage genomes?

Given the above analyses for selecting appropriate thresholds to identify ARG in reference genome datasets, we next applied the conservative and exploratory thresholds to 25 publicly-available human-or animal-associated viromes. These include 10 from the human lung with and without cystic fibrosis (Willner et al 2009), 9 from human feces of healthy subjects (Kim et al 2011, Minot et al 2011), 4 from mouse feces with and without antibiotic treatment (Modi et al 2013), as well as 2 additional viromes from human feces (Minot et al 2012, Reyes et al 2010) where only assembled contigs are available.

Comparison of the virome reads against ARDB+ with a conservative threshold gave no ARG hit for all human feces viromes, a single hit for the ciprofloxacin-treated sample of mice viromes, and 25 hits for the lung samples (Supplementary information S8). Application of an exploratory cut-off (BLASTx, bit-score>70) yielded 0% - 0.19% of virome reads as possible ARG, depending upon the virome, with the lung samples again outliers (Supplementary information S8). Therefore, at this stage, ARG did not appear more abundant in viromes than in viruses, except possibly in lung samples.

### Variable yet often non-negligible presence of bacterial DNA in viromes

Bacterial DNA is common in viromes (Roux et al 2013) and can derive either from phage-encoded bacterial genes (this is what we are looking for in the present study), generalized transduction (a well known process whereby a bacterial DNA segment of the size of the viral genome is taken mistakenly into the capsid instead of the viral DNA), or from insufficient removal of bacterial DNA (either free or entrapped into vesicles) prior phage particle DNA purification. Discriminating between these possibilities is critical towards understanding the mobility of the bacterial DNA detected in the viromes. Currently, when >0.2 ‰ of reads match to 16S ribosomal DNA, a non-viral marker of bacterial DNA, the viromes are considered to have high bacterial DNA content (Roux et al 2013), suggestive of external contamination. Examination of the reads from the 23 unassembled viromes for 16S ribosomal DNA content showed that 17 of the viromes had low (<0.2‰ of 16S DNA reads) levels of bacterial DNA content, whereas 6 lung samples had high levels (Figure 3A, upper panel).

**Figure 3:**
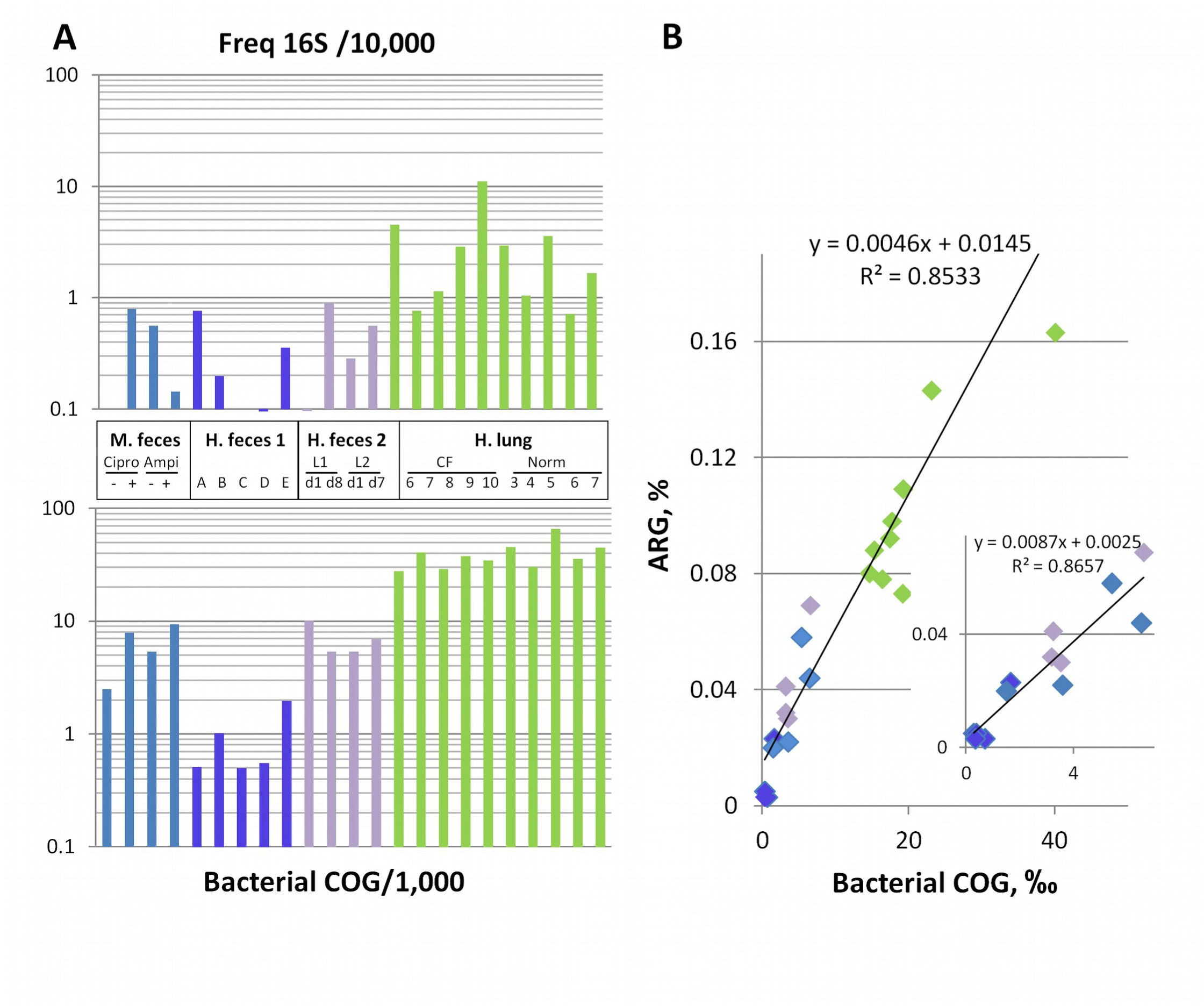
A. Levels of bacterial DNA content in the 23 unassembled viromes. “M. feces” mouse feces samples from Modi et al (2013), Amp, ampicillin, Cipro, ciprofloxacin, mice treated +/- with without antibiotic. “H. feces 1” Human feces samples from Kim et al (2011). “H feces 2” Human feces viromes from Minot et al (2011), L1 and L2 are two lean subjects, at two different time points. “H. lung” Human lung samples from Willner et al (2009), CF cystic fibrosis patients, Norm, healthy subjects. **Upper panel**. Proportion of reads matching against 16S DNA. **Lower panel**. Proportion of reads matching against bacteria-only COGs. **B. Correlation between bacterial (x-axis) and ARG DNA (y-axis) amounts in human-associated viromes**. Data are taken from Supplementary information S8, ARG matches are with the exploratory threshold, bit-score >70. The same color code as in Panel A is applied. Inset: same plot after removal of the lung sample data points.

To increase sensitivity relative to the single 16S gene approach, we established “bacteria-only” protein families by screening the 1,112 most frequently observed COGs in gut microbiomes (Qin et al 2010) against viral proteins derived from NCBI RefSeq genomes. This revealed 688 COGs with no similarities to viral proteins, which we termed “bacteria-only COGs”. The virome reads were then compared to bacteria-only COGs, which largely corroborated the findings from the 16S analyses (Figure 3A, lower panel), but revealed that all 10 lung samples had similar high levels of bacterial DNA (1.5-4% of matches to bacteria-only COGs).

### The more bacterial DNA, the more ARG detected

Previously, the mouse feces viromes (Modi et al 2013) were used to examine the impact of 8-week antibiotic treatments (ampicillin or ciprofloxacin) on the frequency and spread of ARG. The major findings were that treated mice contained ^~^3-fold increase in reads identified as ARG compared to untreated animals, which was interpreted to suggest that ARG were spread through phage genomes. Notably, however, our re-analyses of these data show that there is also a 2- to 3-fold increase in the bacteria-only COGs across these same treatments (Figure 3A, lower panel, blue bars). This suggests that all types of bacterial genes are more frequently detected in the treated mouse viromes, with no particular selection for ARG. Mechanistically, this may be due to the antibiotic-treatment inducing prophages, with some subset performing generalized transduction.

Further, the bacteria-only COGs help uncover background bacterial DNA contamination that confounds interpretations of ARG frequencies across all 23 viromes examined here. Specifically, the percentage of reads matching ARG using the exploratory cut-off (bit-score>70, against ARDB+) is correlated to that for bacteria-only COGs for all 23 viromes (Figure 3B). The fact that the quantities of ARG and bacterial DNA are correlated suggests that the ARG signal in viromes derives from bacterial rather than phage genomes.

### ARG are rare in viral contigs

At this point it appears that the majority of the ARG in viromes are confounded with regular bacterial genes, due to the exploratory threshold used in virome publications (Fancello et al 2011, Modi et al 2013). However, there may still be some cases of ARG in viruses, as suggested in the mice study, and hypothesized to be due to long-term high dose antibiotic treatment (Modi et al 2013).

To separate the bacterial from the viral signal, and gain further insight into whether phages from viromes encoded ARG in their genomes, we assembled the viromes and examined the resulting contigs for the genomic context of ARG. While this will only evaluate the “dominant” viral and transducing DNA of the samples (20% of all reads for 454 viromes map to >2kb-long contigs), it offers yet another opportunity to assess the origin of ARG in viromes. In a first step, all contigs of a size above 2kb were assigned to one of the three following categories: (i) *viral* when a majority of genes matched to viral proteins or proteins of unknown function (with a minimum of 1 viral gene), (ii) *bacterial* when a majority of genes matched bacterial genes and had no viral-typical genes, or (iii) *unknown* where too few genes (<3), a majority of genes of unknown function, or as many bacterial and viral genes (+/-20%) resided on the contig.

Next, contigs of each category were searched for the presence of ARG, using BLAST bit-score>70 and Resfams cut-offs as exploratory and conservative thresholds, respectively. Again, such analyses revealed them to be very uncommon (summarized in Table 1, and Supplementary information S9). Among 465 contigs from mouse samples classified as viral, not a single one contained an ARG, even with ARG identified using the exploratory cut-off. In the lung viromes, as expected from the read analysis, most (68%) of the contigs were bacterial, and ARG represented 0.2% (conservative) to 3.5% (exploratory) of the total genes on these bacterial contigs. None of the lung contigs of viral origin contained ARG. Globally across all viral contigs from all 25 viromes, not a single ARG was detected with the conservative cut-off, and only 5 using the exploratory cut-off (Table 1, map of the contigs Supplementary information S10). All 5 were putative beta-lactamases, and belonged to the same Pfam family as the Palmer-encoded metal-hydrolase tested above, which did not degrade beta-lactams. Still, in some *Bacteroides* prophages, a gene of this same family has reported antibiotic resistance activity (Ogilvie et al 2013), so that experimental testing is needed to validate the prediction.

**Table 1:**
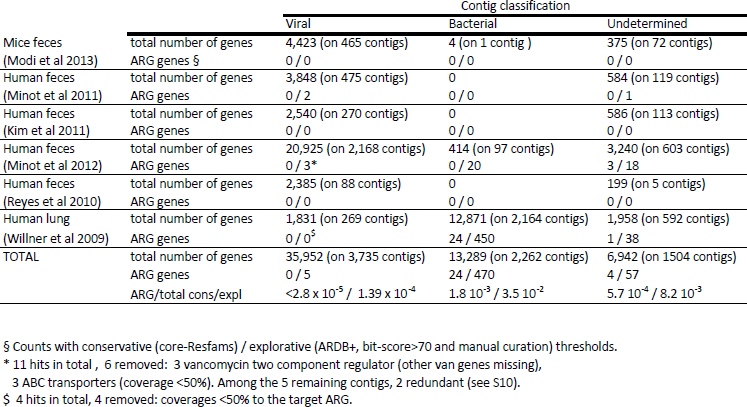
ARG detected on viral and bacterial contigs >2kb, among the 25 viromes

Thus among 35,952 proteins predicted across all available human-and mouse-associated contigs of viral origin, a total of 0 (conservative) and 5 (exploratory) putative ARG were found which suggests an *in silico* frequency of <2.8 × 10^−5^ (conservative) and 1.4x10^−4^ (exploratory) for phage-encoded ARG. In sequenced phages, 1 and 13 putative ARG were detected *in silico*, with the same cut-offs, for 121,506 total proteins, amounting to similar ratio of 0.8x10^−5^ and 1.1x10^−4^. We conclude that frequencies of phage-encoded ARG in viromes are no higher than those in sequenced phage genomes.

After completion of this analysis, a study focusing on saliva and feces virome samples of human subjects that had been treated or not with antibiotics was published (Abeles et al 2015). ARG frequency in virome samples was in the 1% range, using a conservative threshold (E-value<10^−30^ at the DNA level, against CARD) and not significantly different between treated and untreated samples (Abeles et al 2015). Such values being 100-fold higher than in the present study, we included these new viromes in a simplified analysis as follows: (i) bacterial contamination level among raw reads was measured with bacteria-only COGs. Feces samples had generally low levels of matches to such COGs (0.68 +/-0.76 %), but saliva sample levels of bacterial DNA were high (similar to lung virome samples, 4.25 +/- 0.87 %) despite a very low 16S content (see Supplementary information S8). (ii) contigs were assembled, ARG were searched for with the Resfams database (conservative prediction), and origin of ARG-positive contigs was determined (see Supplementary information S9). Among all feces viromes, ten contigs had a putative ARG, two of which were of viral origin. Dividing by the total number of genes found on all feces contigs gave an overall frequency of 7x10^−5^. We conclude that the ARG frequency in this dataset is in a range similar to other viromes. The prior over-estimation might be related to the predominance of the CARD category ‘drug transporters’ (Abeles et al 2015). Our conservative analysis used Core-Resfams, in which some transporters have been excluded, due to their large promiscuity (Gibson et al 2015).

## Discussion

As sequence analysis of uncultivated viral communities becomes more widespread, we present these analyses as a case study to help establish the boundaries of viromic inference. Earlier work has suggested ARG were enriched in the genomes of antibiotic-treated phage communities, both in the lungs of cystic fibrosis patients (Fancello et al 2011) and in the feces of mice (Modi et al 2013). However, reanalysis of these data suggests that the lung sample conclusions were misled by excessive bacterial DNA content, and the mice virome analyses suffered from inflated false positives due to relaxed thresholds for *in silico* detection of ARG (E-value <10^−3^ on ARDB+). To guide future work, we suggest that (i) bacterial DNA contamination be quantified in viromes using analyses such as those presented here and/or automated software now available (VirSorter, Roux et al 2015), (ii) automated analyses use a conservative threshold to quantify *bona fide* ARG, and (iii) discovery-based work proceed with added caution. Specifically, the latter exploratory cut-offs should utilize a bit-score >70 threshold complemented with manual inspection for removing the kind of false positives identified in this study. Additionally, assembly in contigs should be used where possible to confirm that novel ARG are really present on viral contigs and thereby, avoid being misled by general transduction or contaminating bacterial DNA. Finally, only experimental testing will ascertain the function of a predicted ORF as an ARG.

The present analysis of 25 human-and animal-associated viromes does not suggest a paradigm shift is needed with respect to ARG content of phages: phage genomes rarely carry ARG, even under intense selection for antibiotic resistance. It will be interesting in the future to investigate whether this remains true in other environments impacted by antibiotics such as soils. We sought to test experimentally whether some of the newly predicted ARG identified with the exploratory cut-off on complete phage genomes were *bona fide* ARG, and found that none of them were. This suggests that one should stick to conservative cut-offs for asserting ARG presence, or complement predictions made with exploratory cut-offs with experimental data. That no particular increase in ARG frequency was observed in the antibiotic-treated mouse viromes suggests that ARG did not become part of viral genomes in these samples, at least in the dominant phage population for which contigs could be assembled. Independently, on human salivary and feces virome samples, a recent study reached the same conclusion of an absence of ARG enrichment upon treatment (Abeles et al 2015). In a context of general warning and concern about the consequences of rampant exposure to antibiotics, this observation is good news as it constrains the spread of ARG by phages predominantly to mechanisms already well-known, such as generalized transduction. Our observation of a 2-3 fold increase of bacterial DNA content in viromes from mice treated with antibiotics suggests that antibiotics might induce prophages, as already suggested for pig microbiota (Allen et al 2011), with a concomitant increase of generalized transduction, and low frequency ARG transfer.

Because metabolic genes are selected for in aquatic phages (Anantharaman et al 2014, Hurwitz et al 2013, Hurwitz et al 2014, Roux et al 2013, Roux et al 2014), we posit that ARG acquisition must be largely counter-selected in phages. Consistent with this, one study reports an attempt to clone ARG out of an activated sludge viral metagenomic library, with no success (Parsley et al 2010). Moreover, ARG are 10-fold less abundant in phages than prophages (Kleinheinz et al 2014). Mechanistically it is likely that an ARG on a mobile element invading a prophage will inactivate it, so that the prophage will no longer replenish the pool of ‘living’ phages. Consistent with this tenant, there are several prophages carrying ARG for which lytic activity has not been obtained (Billard-Pomares et al 2014, Wipf et al 2014). In particular, three well-studied ARG carrying prophages in *Streptococcus pyogenes* have not been shown yet to be capable of phage lytic activity (Banks et al 2003, Brenciani et al 2010, Iannelli et al 2014). The three elements carry over the antibiotic resistance phenotype across Streptococcal species when cells are put into contact, using a conjugation protocol (Giovanetti et al 2014, Santagati et al 2003). Whether these ARG-carrying prophages that may not be lytically active represent an important category of mobile elements in terms of ARG spreading, remains an open question.

In summary, these re-analyses present a roadmap for drawing robust conclusions about ARG, and more generally all types of bacterial genes in virome datasets. As a result, the emerging picture for the spread of ARG suggests that despite the excessive use of antibiotics in humans and animals, ARG may be among the “bacterial host genes” that have not (yet?) been selected for in phage genomes, at least in the human and mice-associated environments studied here.

## Acknowledgements

We thank Graham Hatfull for the gift of phages Mozy and Corndog, and Gabriel Everett for the gift of phage Palmer. Marie Touchon, Etienne Ruppé and Alexandra Gruss are also thanked for their critical reading of the manuscript.

